# Gut bacterial aggregates as living gels

**DOI:** 10.1101/2021.06.08.447595

**Authors:** Brandon H. Schlomann, Raghuveer Parthasarathy

## Abstract

The spatial organization of gut microbiota influences both microbial abundances and host-microbe interactions, but the underlying rules relating bacterial dynamics to large-scale structure remain unclear. To this end we studied experimentally and theoretically the formation of three-dimensional bacterial clusters, a key parameter controlling susceptibility to intestinal transport and access to the epithelium. Inspired by models of structure formation in soft materials, we sought to understand how the distribution of gut bacterial cluster sizes emerges from bacterial-scale kinetics. Analyzing imaging-derived data on cluster sizes for eight different bacterial strains in the larval zebrafish gut, we find a common family of size distributions that decay approximately as power laws with exponents close to −2, becoming shallower for large clusters in a strain-dependent manner. We show that this type of distribution arises naturally from a Yule-Simons-type process in which bacteria grow within clusters and can escape from them, coupled to an aggregation process that tends to condense the system toward a single massive cluster, reminiscent of gel formation. Together, these results point to the existence of general, biophysical principles governing the spatial organization of the gut microbiome that may be useful for inferring fast-timescale dynamics that are experimentally inaccessible.

## Introduction

The bacteria inhabiting the gastrointestinal tracts of humans and other animals make up some of the densest and most diverse microbial ecosystems on Earth [1, 2]. In both macroe-cological contexts and non-gut microbial ecosystems, spatial organization is well known to impact both intra- and inter-species interactions [3, 4, 5]. This general principle is likely to apply in the intestine as well, and the spatial structure of the gut microbiome is increasingly proposed as an important factor influencing both microbial population dynamics and health-relevant host processes [6, 7]. Moreover, recent work has uncovered strong and specific consequences of spatial organization in the gut, such as proximity of bacteria to the epithelial boundary determining the strength of host-microbe interactions [8, 9], and antibiotic-induced changes in aggregation causing large declines in gut bacterial abundance [10]. Despite its importance, the physical organization of bacteria within the intestine remains poorly understood, in terms of both in vivo data that characterize spatial structure and quantitative models that explain the mechanisms by which structure arises.

Recent advances in the ability to image gut microbial communities in model animals have begun to reveal features of bacterial spatial organization common to multiple host species. Bacteria in the gut exist predominantly in the form of three-dimensional, multicellular aggregates, often encased in mucus, whose sizes can span several orders of magnitude. Such aggregates have been observed in mice [11], fruit flies [12], and zebrafish [13, 14, 15, 10, 9], as well as in human fecal samples [16]. However, an understanding of the processes that generate these structures is lacking.

The statistical distribution of object sizes can provide powerful insights into underlying generative mechanisms, a perspective that has long been applied to datasets as diverse as galaxy cluster sizes, [17], droplet sizes in emulsions [18], allele frequency distributions in population genetics [19], immune receptor repertoires [20], species abundance distributions in ecology [21], protein aggregates within cells [22], and linear chains of bacteria generated by antibody binding [23].

Motivated by these analogies, we sought to understand the distribution of three-dimensional bacterial cluster sizes in the living vertebrate gut, aiming especially to construct a quantitative theory that connects bacterial-scale dynamics to global size distributions. Such a model could be used to infer dynamical information in systems that are not amenable to direct observation, such as the human gut. Identifying key processes that are conserved across animal hosts would further our ability to translate findings in model organisms to human health-related problems. At a finer level, validated mathematical models could be used to infer model parameters of specific bacterial species of interest, for example pathogenic invaders or deliberately-introduced probiotic species, by measuring their cluster size distribution.

We analyzed bacterial cluster sizes obtained from recent imaging-based studies of the larval zebrafish intestine [14, 10, 9]. As detailed below, we find a common family of cluster size distributions with bacterial species-specific features. We show that these distributions arise naturally in a minimal model of bacterial dynamics that is supported by direct observation. The core mechanism of this model involves growth together with a fragmentation process in which single cells leave larger aggregates. Strikingly, this process can be mapped exactly onto population genetics models of mutation, with cluster size analogous to allele frequency and single cell fragmentation analogous to mutation. The combination of growth and fragmentation generates size distributions with power law tails, consistent with the data. Further, we show that cluster aggregation can generate an overabundance of large clusters through a process analogous to the sol-gel transition in polymer and colloidal systems, leading to plateaus in the size distribution that are observed in the data. In summary, we find that gut microbiota can be described mathematically as “living gels”, combining the statistical features of evolutionary dynamics with those of soft materials. Based on the generality of our model and our observations across several different bacterial species, we predict that this family of size distributions is universal across animal hosts, and we provide suggestions for testing this prediction in various systems.

## Results

### Different bacterial species share a common family of broad cluster size distributions in the larval zebrafish intestine

We combined and analyzed previously-generated datasets of gut bacterial cluster sizes in larval zebrafish [14, 10, 9]. In these experiments, zebrafish were reared devoid of any microbes, i.e. “germ-free”, and then mono-associated with a single, fluorescently labelled bacterial strain (Fig. 1A). After a 24 hour colonization period the complete intestines of live hosts were imaged with light sheet fluorescence microscopy [24, 25] (Fig. 1B). Bacteria were identified in the images (Fig. 1C) using a previously described image analysis pipeline [13, 14]. Single bacterial cells and multicellular aggregates were identified separately, and then the number of cells per multicellular aggregate was estimated by dividing the total fluorescence intensity of the aggregate by the mean intensity of single cells (Methods).

**Figure 1:**
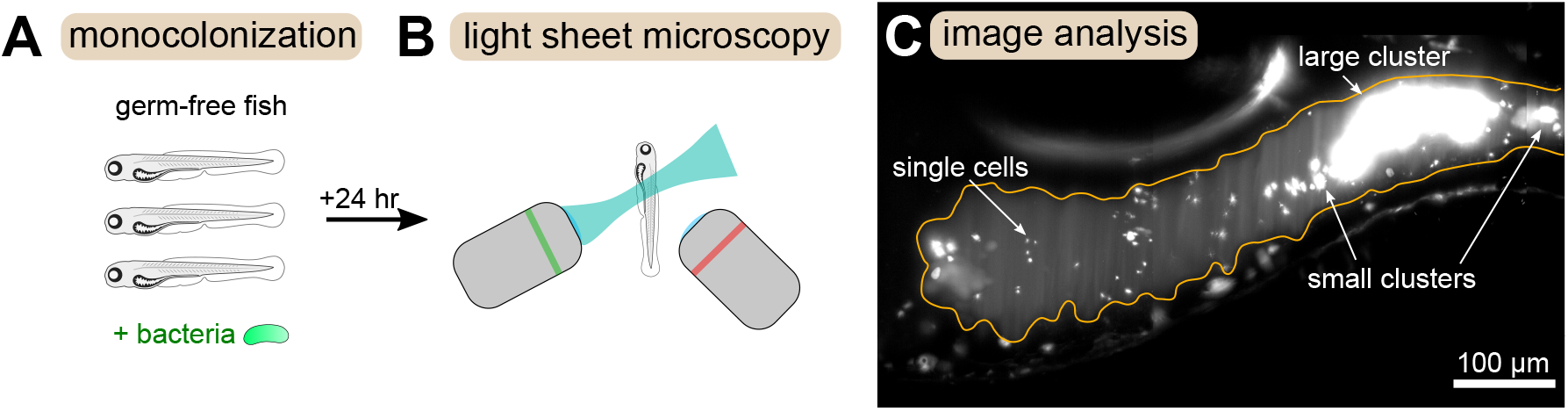
Overview of experimental methods. Larval zebrafish were derived germ-free and then monoassociated with single bacterial species (left). After 24 h of colonization, images spanning the entire gut were acquired with light sheet fluorescence microscopy (middle). An example image of the anterior intestine is shown on the right, with instances of single cells and multicellular aggregates marked. The image is a maximum intensity projection of a 3D image stack. The approximate boundary of the gut is outlined in orange. Sizes of bacterial clusters were estimated with image analysis by separately identifying single cells and multicellular aggregates, and then normalizing the fluorescence intensity of aggregates by the mean single cell fluorescence.

In total, we characterized eight different bacterial strains, summarized in Table 1. Six of the strains were isolated from healthy zebrafish [26] and then engineered to express fluorescent proteins [27], and two are genetically engineered knockout mutants of *Vibrio* ZWU0020, defective in motility (“Δmot”) and chemotaxis (“Δche”), as described in reference [9]. The parent strain of these mutants, *Vibrio* ZWU0020, scarcely forms aggregates at all, existing primarily as single, highly motile cells [15, 10, 9], and so was excluded from this analysis. All strains are of the phylum Proteobacteria [27]. A table of all cluster sizes by sample is included in the Supplementary Data File.

**Table 1:** Summary of cluster data by bacterial strain. Each row corresponds to one of the bacterial strains included in this study. Entries include strain name, total number of fish colonized with that strain, total number of clusters identified across all fish, and the original publication that the data was pulled from.

| bacterial strain | number of fish | number of clusters | source publication |
| --- | --- | --- | --- |
| *Aeromonas* ZOR0001 | 6 | 445 | [14] |
| *Aeromonas* ZOR0002 | 6 | 1901 | [14] |
| *Enterobacter* ZOR0014 | 18 | 3597 | [14, 10] |
| *Plesiomonas* ZOR0011 | 3 | 223 | [14] |
| *Pseudomonas* ZWU0006 | 6 | 133 | [14] |
| *Vibrio* ZOR0036 | 6 | 2430 | [14] |
| *Vibrio* ZWU0020 Δmot | 11 | 5888 | [9] |
| *Vibrio* ZWU0020 Δche | 11 | 3551 | [9] |

We calculated for each bacterial strain the reverse cumulative distribution of cluster sizes, *P*(size > *n*), denoting the probability that an intestinal aggregate will contain more than *n* bacterial cells. We computed *P*(size > *n*) separately for each animal (Fig. 2, small circles) and also pooled the sizes from different animals colonized by the same bacterial strain (Fig. 2, large circles). There is substantial variation across fish, but the pooled distributions exhibit a well-defined average of the individual distributions. We also computed binned probability densities (Fig. S1), which show similar patterns, but focus our discussion on the cumulative distribution to circumvent technical issues related to bin sizes.

**Figure 2:**
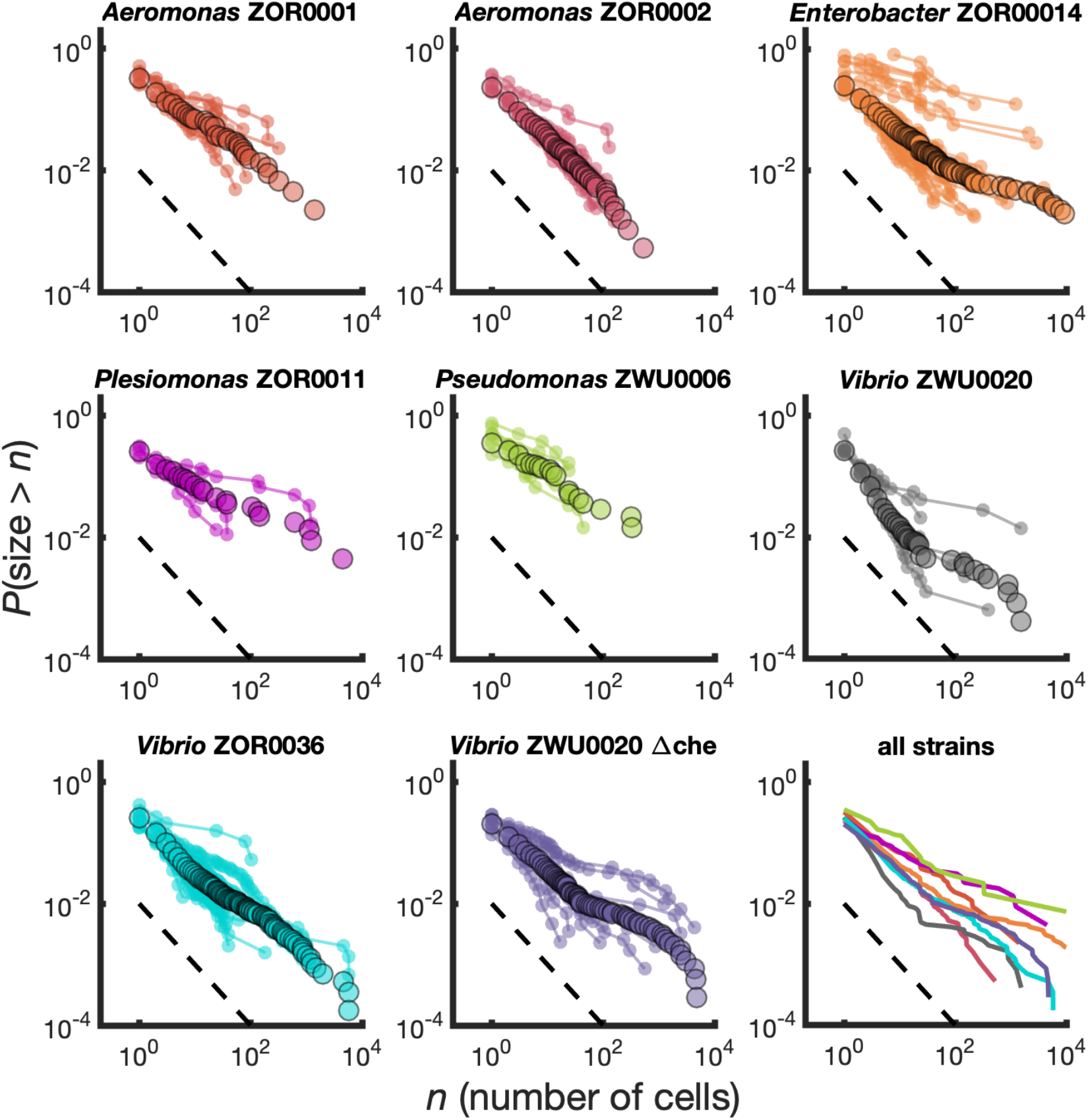
Different bacterial species exhibit similar cluster size distributions. Reverse cumulative distributions, the probability that the cluster size is greater than *n* as a function of *n*, for eight bacterial strains in larval zebrafish intestines. Small circles connected by lines represent the distributions constructed from individual fish. Large circles are from pooled data from all fish. The dashed line represents *P*(size > *n*) ~ *n*^−1^. Bottom right panel shows the pooled distributions for each strain as solid lines.

We find broad distributions of *P*(size > *n*) across all strains (Fig. 2, bottom right panel). For comparison, for each strain we overlay a dashed line representing the power law distribution *P*(size > *n*) ~ *n*^−1^ corresponding to a probability density of *p*(*n*) ~ *n*^−2^. Each strain’s cumulative distribution follows a similar power-law-like decay at low *n*, with an apparent exponent in the vicinity of −1, and then becomes shallower in a strain-dependent manner. For example, *Aeromonas* ZOR0002 has a quite straight distribution on a log-log plot (Fig. 2, top row, middle column), while the distribution of *Enterobacter* ZOR0014 exhibits a plateau-like feature at large sizes (Fig. 2, top row, right column). The mutant strains *Vibrio* ZWU0020 Δche and Δmot follow qualitatively similar distributions to the native strains (Fig. 2, bottom row, left and middle columns).

In summary, we find that different bacterial strains, which exhibit a variety of swimming and sticking behaviors [27, 14], abundances [14, 10, 9, 15], and population dynamics [15, 10, 9], share a common family of cluster size distributions. This observation suggests that generic processes, rather than strain-specific ones, determine gut bacterial cluster sizes. Notably, these distributions are extremely broad, inconsistent with the exponential-tailed distributions found for linear chains of bacteria [23]. We next sought to understand the kinetics that give rise to our measured cluster size distributions.

### A growth-fragmentation process generates power-law distributions

Previous time-lapse imaging of bacteria in the zebrafish intestine revealed four core processes that can alter bacterial cluster sizes: (1) clusters can increase in size due to cell division, a process we refer to as “growth”; (2) clusters can decrease in size as single bacteria escape from them, a process we refer to as “fragmentation” and believe to be linked to cell division at the surface; (3) clusters can increase in size by joining with another cluster during intestinal mixing, a process we refer to as “aggregation”; and (4) clusters can be removed from the system by transiting along and out of the intestine, a process we refer to as “expulsion”. We previously showed that a mean-field kinetic model containing these processes and parameterized by independent experiments generates a size distribution consistent with that of *Enterobacter* ZOR0014 [10]. However, it was not clear which processes generated which features of the distribution, or how generalizable the model was. Therefore, we studied this model in more detail, starting from a simplified version and iteratively adding complexity.

The observation that all distributions appeared to be organized around *P*(size > *n*) ~ *n*^−1^ inspired us to consider connections to a classic populations genetics model that has this form for the distribution of allele frequencies, known as the Yule-Simons process [28, 29, 30, 19]. An exponentially growing population subject to random neutral mutations that occur with probability *ϵ* will amass an allele frequency distribution that follows 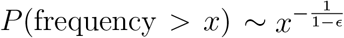 for large sizes, with the limit to *P*(frequency > *x*) ~ *x*^−1^ for rare mutation. This heavy-tailed distribution reflects “jackpot” events in which mutants that appear early rise to large frequencies through exponential growth.

Analogously, the size of mutant clones maps onto the size of a bacterial cluster, and the mutation process that generates new clones maps onto the fragmentation process that generates new clusters (Fig. 3A). In situations where all cells in a cluster have the same probability of fragmenting, this analogy is exact and the same distribution emerges (Supplementary Text). However, gut bacterial clusters are three-dimensional and likely encased in mucus [16], so spatial structure likely influences fragmentation rates. We hypothesized that this spatial structure could be a mechanism for generating distributions shallower than *P*(size > *n*) ~ *n*^−1^ that we observe in the data for large sizes (Fig. 2) but that cannot be produced by the standard Yule-Simons mechanism. Therefore, we modified the Yule-Simons process by decoupling the growth and fragmentation processes and invoking a fragmentation rate, *F_n_*, that scales as a power of the cluster size, *F_n_* ~ *βn^ν_F_^* (Fig. 3B). A value of *ν_F_* = 1 corresponds to the well-mixed limit of the Yule-Simons process. A value of *ν_F_* = 2/3 corresponds to only bacteria on the surface of clusters being able to fragment. An extreme value of *ν_F_* = 0 means that all clusters have the same rate of fragmenting, regardless of their size, and can be thought of as representing a chain of cells where only the cells at ends of the chain can break off.

**Figure 3:**
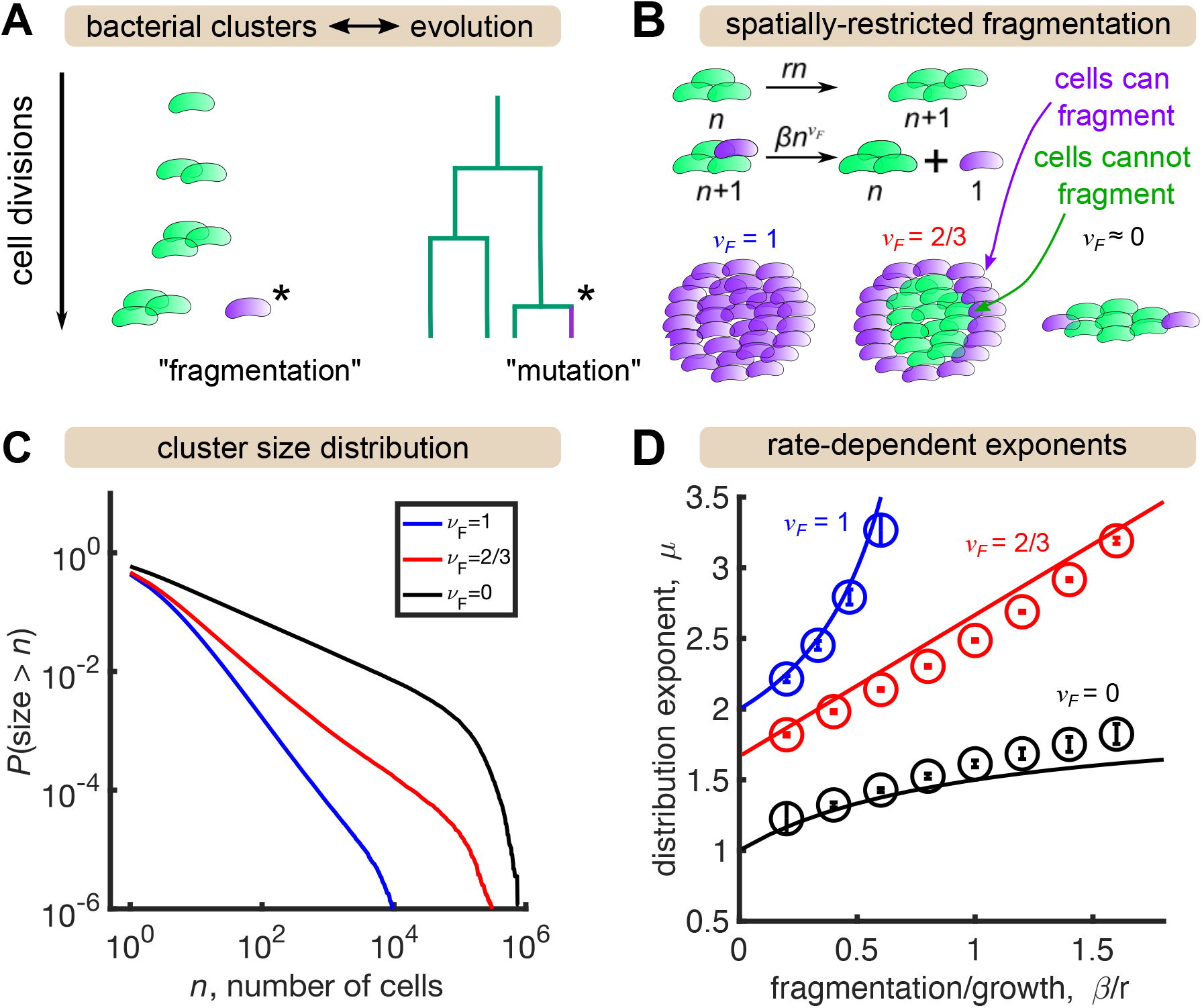
A growth and fragmentation process maps to evolutionary dynamics and generates power law distributions. (A) Fragmentation is analogous to mutation and we can construct a genealogy that mirrors the physical structure of the clusters. (B) Summary of a growth/fragmentation process that includes the effect of spatially-confined clusters. (C) Examples of reverse cumulative size distributions obtained from stochastic simulations of the model for different values of the fragmentation exponent, *ν_F_*. The tails of the distribution are approximately power laws, defined as *P*(size > *n*) ~ *n*^−*μ*+1^. Parameters: *r* = 0.5 hr^−1^, *β* = 0.4, 0.2, 0.167 hr^−1^, for *ν_F_* = 0, 2/3, 1, respectively, time *t* = 24 hr, and the system was initialized with 10 single cells. (D) Dependence of the resulting distribution exponent, *μ*, on ratio of fragmentation to aggregation rate (*β/r*) and fragmentation exponent (*ν_F_*). Markers show mean and standard deviation across 100 simulations. Solid lines are approximate analytic results (Table 2). Parameters: same as (C) with *β* varying

In stochastic simulations of this model (Methods) we find broad, power-law-like distributions for each value of *ν_F_* (Fig. 3C), but no signature of a shallow plateau at larger sizes. Following established methods, we fit a power law, *P*(size > *n*) ~ *n*^−*μ*+1^ for *n* > *n*_min_ to simulation outputs using maximum likelihood estimation [31] and examined the dependence on fragmentation rate. Faster fragmentation results in larger values of *μ*, reflecting steeper distributions, with the dependence being superlinear for *ν_F_* = 1, approximately linear for *ν_F_* = 2/3, and sublinear for *ν_F_* = 0 (Fig. 3D, circles). Increasing values of *ν_F_* also appeared to have increasing minimum values of *μ*, corresponding to rare fragmentation.

The minimum value of the distribution exponent can be computed by considering, for example, the total rate of fragmentation events. Denoting the total number of clusters by *M* and the number of clusters of size *n* by *c_n_*, the rate of cluster production follows *M* ≈ *β*Σ_*n*_ *n^ν_F_^ C_n_* (Supplementary Text). Assuming a power-law solution *c_n_* ~ *n*^−*μ*^ and approximating the sum by an integral, we see that the rate of cluster production is finite only if

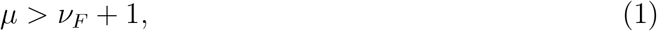

consistent with simulations. Therefore, spatial structure—modeled by decreasing *ν_F_*—is indeed a mechanism to generate distributions shallower than *P*(size > *n*) ~ *n*^−1^. A heuristic argument for the rate dependence of the exponents in the long time, large size limit is provided in the Appendix, with the results summarized in Table 2 and plotted as solid lines in Fig. 3D. The analytic results agree reasonably well with simulations, with deviations becoming prominent as *β/r* ≈ 1.

**Table 2:** Analytic results for the minimal growth-fragmentation process. Distribution exponent, *μ*, as a function of fragmentation exponent, *ν_F_*, fragmentation rate, *β*, and growth rate, *r*, as plotted in Fig. 3D. Results are expected to be valid for long times (*t* → ∞), large sizes (*n* → ∞), and slow fragmentation (*β/r* < 1). See Supplementary Text for details.

| | $\nu_F = 1$ | $\nu_F = 2/3$ | $\nu_F = 0$ |
| --- | --- | --- | --- |
| distribution exponent, $\mu$ | $1 + \frac{1}{1-\beta/r}$ | $\frac{5}{3} + \frac{\beta}{r}$ | $1 + \frac{\beta/r}{1+\beta/r}$ |

In summary, we identified a minimal growth-fragmentation process that generates power-law distributions with tuneable exponents in the experimentally observed range. However, this model does not include other features known to occur in the experimental system, including a finite carrying capacity that limits growth, cluster aggregation, and cluster expulsion, which may alter the asymptotic distributions. Moreover, this model fails to capture the large-size behavior of many of the experimental distributions, which exhibit a shallowing or plateau effect (Fig. 2). Therefore, we investigated extensions of the model.

### Size-dependent aggregation enhances the abundance of large clusters

We explored a number of potential mechanisms for generating plateaus in the size distribution at large cluster sizes. As shown below, several plausible models fail to produce this feature. It emerges, however, from the incorporation of size-dependent aggregation rates.

First we tested whether finite time effects could introduce plateaus to the distributions of the minimal growth-fragmentation model, since our power-law solutions are only valid asymptotically. Indeed, stochastic simulations with *ν_F_* = 1 and rare fragmentation (*r* = 0.5 hr^−1^, *β* = 0.05 hr^−1^) showed that for systems initialized with 10 single cells (a reasonable comparison with initial colonization in the experiments [15]), slight curvature appears in the distribution that weakens with time but is still detectable at 24 hours (Fig. S2, circles). We confirmed that this effect was solely due to dynamics and not to any finite system effect by numerically integrating the master equation for this model, which describes the deterministic dynamics of an infinite system yet agrees with the stochastic simulation results (Fig. S2, lines; Methods). However, the curvature observed at finite times is substantially smaller than what occurs for some of the strains, such as *Enterobacter* ZOR0014 and *Vibrio* ZOR0036, so we believe it is not the dominant effect.

We next asked whether including additional processes to the model could produce the plateau effect, focusing on stationary distributions. As discussed above, populations in the larval zebrafish gut are known to reach carrying capacities that halt growth [13]. Since we believe fragmentation is tied to growth, we modeled this as the fragmentation rate being slowed as the total number of cells, *N*, approaches carrying capacity, *K*, in the same way as the growth rate: *r* → *r*(1 — *N/K*) and *β* → *β*(1 — *N/K*). Carrying capacities have been estimated to range from 10^3^ — 10^6^ cells, depending on the bacterial strain [13, 15, 14, 10, 9].

With this addition to the model, fragmentation halts in the steady state. However, in the larval zebrafish gut is has been well-documented that large bacterial aggregates are quasi-stochastically expelled out the intestine, after which exponential growth by the remaining cells is restarted [15, 9]. We modeled expulsion by having clusters removed from the system altogether at a size-dependent rate *E_n_* = λ*n^ν_E_^*. It is unclear what value of the exponent *ν_E_* best describes the experimental system, but previous studies measured expulsion rates for the largest clusters, typically of order *K* ~ 10^3^ cells, in the range of 0.07 to 0.11 hr^−1^ [15, 9, 10]. Therefore, we co-varied *ν_E_* and λ such that λ*K^ν_E_^*~ 10^−1^ hr^−1^. Combining finite carrying capacity and expulsion leads to a non-trivial stationary distribution of the model that lacks a plateau for *ν_E_* = 0, 1/3, or 2/3 (Fig. S3).

Finally, we considered the effect of cluster aggregation, which has been directly observed in live imaging experiments [10]. We model aggregation with pairwise interactions where clusters come together and form a single cluster with size equal to the sum of the individual sizes. The aggregation rate is allowed to be size-dependent with the homogenous kernel *A_nm_* = *α*(*nm*)^*ν_A_*^. As with expulsion, it is not clear which exponent value is the most realistic. Accurate measurements of aggregation rates are lacking, but we estimate bounds to be between 1 and 100 total aggregation events per hour for a typical population (Methods), so we consider only pairs of *α* and *ν_A_* that match these bounds. Further, an important theoretical distinction is that in purely aggregating systems, models with *ν_A_* ≥ 1/2 exhibit a finite-time singularity corresponding to a gelation transition, at which point the distribution acquires a power-law tail, while distributions have exponential tails when *ν_A_* < 1/2 [32]. We considered both regimes.

We added aggregation to our growth-driven process and arrived at the general model described in Fig. 4A. Parameters are also summarized in Table 3. Strikingly, we found that increasing aggregation rate produces the large-size plateau seen in our data, but only when aggregation rate scales sufficiently quickly with cluster size (Fig. 4B, right, *ν_A_* = 2/3) and not when aggregation is size-independent (Fig. 4B, left, *ν_A_* = 0). A mild effect is observed for *ν_A_* = 1/3 (Fig. 4B, middle). The largest plateau (Fig. 4B, *ν_A_* = 2/3, highest curve) corresponds to 15 ± 3 (mean ± std. dev) total aggregation events per hour. This value is consistent with our rough experimental bounds of 1-100 hr^−1^.

**Figure 4:**
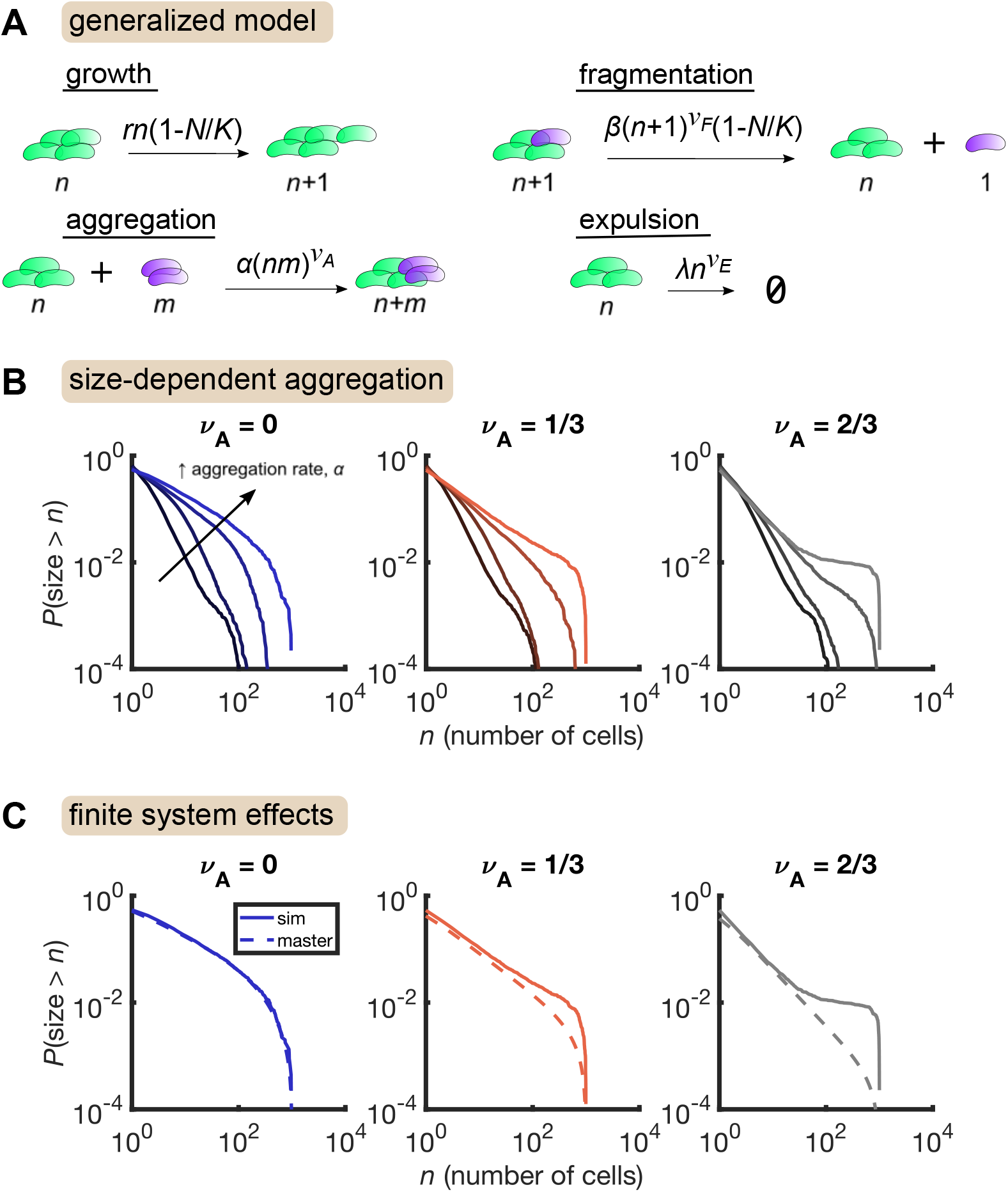
Size-dependent aggregation introduces a plateau in the size distribution. (A) Schematic of the generalized model. Parameters summarized in Table 3. (B) Reverse cumulative distributions obtained from simulations for different values of *ν_A_* (left, middle, right) and *α* (different colored lines within each panel). Increasing aggregation produces a plateau if the aggregation depends strongly enough on cluster size. (C) The plateau arises only in stochastic simulation of finite systems with size-dependent aggregation. Solid lines are stochastic simulations, dashed lines are the result of numerically integrating the master equation. Parameters: *r* = 0.5 hr^−1^, *ν_F_* = 2/3, *β* = 0.5 hr^−1^, *ν_E_* = 1/3, λ = 0.01 hr^−1^, *K* = 10^3^, and the number of simulation was replicates = 150 per parameter set. For each value of *ν_A_*, we considered *α* values of 0 (no aggregation) and then varied *α* logarithmically, with the following (min, max) values for log_10_ *α*: (−4,−2) for *ν_A_* = 0, (−4.5,−2.5) for *ν_A_* = 1/3, and (−5,−3) for *ν_A_* = 2/3.

**Table 3:** Summary of model parameters.

| parameter | description |
| --- | --- |
| $r$ | cell division rate |
| $K$ | carrying capacity |
| $\beta$ | fragmentation rate |
| $\nu_F$ | fragmentation exponent |
| $\alpha$ | aggregation rate |
| $\nu_A$ | aggregation exponent |
| $\lambda$ | expulsion rate |
| $\nu_E$ | expulsion exponent |

We further found that this plateau effect is intrinsic to finite systems (Fig. 4C). For the most aggregated cases in Fig. 4B, we numerically solved the corresponding master equation, representing the deterministic dynamics of an infinite system, and found that the plateau did not occur. Master equation and stochastic simulation solutions agree for *ν_A_* = 0, but for larger values of *ν_A_*, the two solutions only agree in the small size regime. At large sizes, stochastic simulations produce an overabundance of large clusters compared to the master equation solution. This result indicates that in a finite system, strong aggregation can deplete small clusters, condensing them into a small number of large clusters on the order of the system-size.

This overall process is reminiscent of the gelation transition in soft materials. Stochastic dynamics of finite systems of purely aggregating particles at the gelation transition also produces distributions with plateaus, but with an initial decay given approximately by a power law with *μ* = 5/2 (Fig. S4, see also [33]). Combined with a growth/fragmentation/expulsion process, we found that size-dependent aggregation produces a distribution that initially decays in a power-law-like manner with tunable exponents and then exhibits a tuneable plateau, as we observe in the experimental data.

## Discussion

We analyzed image-derived measurements of bacterial cluster sizes from larval zebrafish intestines and discovered a common family of size distributions shared across bacterial species. These distributions are extremely broad, exhibiting a power-law-like decay at small sizes that becomes shallower at large sizes in a strain-specific manner. We then demonstrated how these distributions emerge naturally from realistic kinetics: growth and single-cell fragmentation together generate power-law distributions, analogous to the distribution of neutral alleles in expanding populations, while size-dependent aggregation leads to a plateau representing the depletion of mid-sized clusters in favor for a single large one. In summary, we found that gut bacterial clusters are well-described by a model that combines the features of evolutionary dynamics in growing populations with those of inanimate systems of aggregating particles; intestinal bacteria form a “living gel”.

Given the general and minimal nature of the model’s assumptions, we predict that the form of the cluster size distributions we described here is common to the intestines of animals, including humans. This prediction of generality could be tested in a variety of systems using existing methods. In fruit flies, live imaging protocols have been developed that have revealed the presence of three-dimensional gut bacterial clusters highly reminiscent of what we observe in zebrafish, particularly in the midgut [12]. Quantifying the sizes of these clusters would allow further tests of our model.

In mice, substantial progress has been made in imaging histological slices of the intestine with the luminal contents preserved [34]. Intestinal contents are very dense in the distal mouse colon, however, and it is not clear how one should define cluster size. Other intestinal regions are likely more amenable to cluster analysis. Moreover, with species-specific labelling, it is possible to measure the distribution of clonal regions in these dense areas [35]. One could imagine then comparing these data to a spatially-explicit, multispecies extension of the model we studied here.

Our model could also be tested indirectly for humans and other animals incompatible with direct imaging by way of fecal samples. Two decades ago, bacterial clusters spanning three orders of magnitude in volume were observed in gently-dissociated fecal samples stained for mucus, but precise quantification of size statistics was not reported [16]. Repeating these measurements with quantification, from for example imaging or flow cytometry, would also provide a test of our model, albeit on the microbiome as a whole rather than a single species at a time. The interpretation therefore would be of an effective species with kinetic rates representing average rates of different species.

To close, we emphasize that the degree of bacterial clustering in the gut is an important parameter for both microbial population dynamics and host-bacteria interactions. More aggregation leads to larger fluctuations in abundance due to the expulsion of big clusters, and also thereby increase the likelihood of extinction [10, 36]. Further, aggregation within the intestinal lumen can reduce access to the epithelium and reduce pro-inflammatory signalling [9]. Therefore, measurements of cluster sizes may be an important biomarker for microbiota-related health issues, and inference of dynamics from size statistics using models like this one could aid the development of therapeutics.

## Methods

### Data

We assembled data on gut bacterial cluster sizes from three different studies on larval zebrafish [14, 10, 9]. Size data from [14] and [10] were taken directly from the supplementary data files associated with those publications. The raw size data from [9] was not included in its associated supplementary data file, but summary statistics such as planktonic fraction were. All sizes were rounded up to the nearest integer.

Details of experimental procedures can be found in the original papers. In brief, as described in Fig. 1, animals were reared germ-free, mono-associated with a single bacterial strain, and then imaged 24 hours later using a custom-built light sheet fluorescence microscope [13]. The gut is imaged in 4 tiled sub-regions that are registered via cross-correlation and manual adjustment. Imaging a full gut volume (≈ 1200 *μ*m×300 *μ*m× 150 *μ*m) with 1 *μ*m slices takes approximately 45 seconds. Laser power (5 mW) and exposure time (30 ms) were identical for all experiments.

The image analysis pipeline used to enumerate bacterial cluster sizes is also described in detail in the original publications and in reference [13]. In brief, single cells (small objects) and multicellular aggregates (large objects) are identified separately. The number of cells per aggregate is then estimated as the total fluorescence intensity of the aggregate divided by the mean fluorescence intensity of a single cell. Small objects are identified in three dimensions with a combination of difference-of-gaussians and wavelet filters [37] and then culled using a support vector machine classifier and manual curation. Large objects are segmented in maximum intensity projections using a graph-cut algorithm [38] seeded by either an intensity-or gradient-thresholded mask. The total intensity of an aggregate is computed by extending the two-dimensional mask in the *z*-direction and summing fluorescence intensities above a threshold calculated from the boundary of the mask, with pixels detected as part of single cells removed. The boundary of the gut is manually outlined prior to image analysis and used to exclude extra-intestinal fluorescence.

### Size distribution

For the experimental data, reverse cumulative distributions were computed as

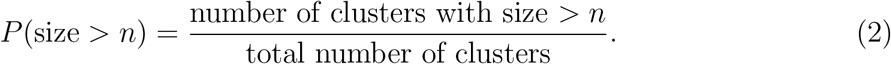

In combining data from different samples colonized with the same strain, we pooled together all sizes and computed the distribution in the same way. For simulations with large numbers clusters, we computed this distribution iteratively, looping through each simulation replicate and independently updating (number clusters with size > *n*) and (total number of clusters), and normalizing at the end.

For the binned probability densities in Fig. S1, data were similarly pooled across samples and then sorted into logarithmically spaced bins of log_10_ width = 0.4.

### Estimates on bounds of agg rates

We estimated approximate bounds on the rate of total aggregation events as follows. For the maximum rate, we note that a typical population contains approximately 200 clusters (mean ± std. dev of 244 ± 182). In the absence of other processes, condensing this system into one cluster would require 100 aggregation events. Populations consisting of almost entirely one large cluster are rare but have been documented [14, 10]. Therefore, we estimate that this complete condensation can occur no more than once an hour, leading to an upper bound on the total rate of aggregation events of 100 per hour.

For the minimum rate, we start with the observation that aggregation has been directly observed between small clusters and also between small clusters and a single large cluster during a large expulsion event [10]. Considering just the latter process, we know that large expulsion events happen roughly once every 10 hours. If approximately 10 small clusters are grouped into the large cluster during transit out of the gut, that would correspond 10 total aggregation events in 10 hours, or, 1 per hour, which we take as a lower bound.

### Simulations

We used three different numerical approaches for studying the models discussed here. The minimal growth-fragmentation process in Fig. 3 was simulated with a Poisson tau-leaping algorithm [39] with a simple fixed tau value of *τ* = 0.1 hr. At each time step the number of growth and fragmentation events was drawn from a Poisson distribution with the rates given in Fig. 3B along with the constraint that clusters must be of size two or larger to fragment.

For the full model including aggregation and expulsion, we used Gillespie’s algorithm [40] for fragmentation, aggregation, and expulsion events, while growth was updated deterministically according to a continuous logistic growth law approximated by an Euler step with *dt* = min(*τ*, 0.1 hr), where *τ* here refers to the time to next reaction. For the Gillespie steps, if the time to next reaction exceeded the doubling time, (ln2)/*r*, the growth steps were performed and then the propensity functions were re-calculated.

Finally, we compared these stochastic simulations to a model in the thermodynamic limit where individual clusters are replaced with cluster densities that evolve deterministically, which is referred to as a master equation [32]. The master equation for the general model reads

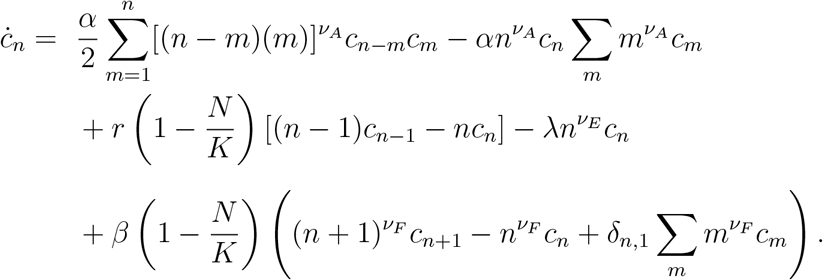

This set of equations was solved numerically on a bounded size grid using an Euler method with step size *dt* = 0.0001 hr. Models that include a carrying capacity, *K*, are already defined on a finite domain of integers ranging from 1 to *K* and the master equation is naturally represented by a set of *K* ordinary differential equations. For models without a carrying capacity, we introduced a maximum size given by the average population size at the last time point, *n*_max_ = exp(*rt*_max_) (rounded up to the nearest integer), and used reflecting boundary conditions at *n*_max_.

A distribution was deemed stationary if it was visibly unchanged after an additional 50% of simulation time.

MATLAB code for simulating these models and plotting data can be found at https://github.com/rplab/cluster_kinetics.

### Estimating distribution exponents

For the simulated distributions in Fig. 3 we estimated a power law exponent using the maximum likelihood-based method described in [31] and the `plfit.m` code supplied therein. This model includes a minimum size as a free parameter that dictates when the power-law tail begins. The minimum size is chosen to minimize the Kolmogorov-Smirnov distance between the data and model distributions for sizes greater than the minimum size. Best fit values of the exponent and minimum size are included in the Supplementary Data File.

## Acknowledgements

We thank Jayson Paulose for helpful discussions. Research was supported by the National Institutes of Health (http://www.nih.gov/), under Awards P50GM09891, P01GM125576, F32AI112094, and T32GM007759. Work was also supported by the National Science Foundation under Award 1427957, and an award from the Kavli Microbiome Ideas Challenge, a project led by the American Society for Microbiology in partnership with the American Chemical Society and the American Physical Society and supported by The Kavli Foundation. The University of Oregon Zebrafish Facility is supported by a grant from the National Institute of Child Health and Human Development (P01HD22486). BHS was supported by the James S. McDonnell Foundation postdoctoral fellowship. The funders had no role in study design, data collection and analysis, decision to publish, or preparation of the manuscript.

## Supplementary Text

### Analytic calculations for growth-fragmentation processes

We consider a model with only growth and fragmentation processes and make heuristic arguments for the form of the asymptotic size distribution. In particular, we are interested in how the exponent of the resulting power law tails depends on the growth and fragmentation rates. We find the following results:

**Table S1:** Analytic results for distribution exponent, *μ*, as a function of fragmentation exponent, *ν_F_*, fragmentation rate, *β*, and growth rate, *r*. Results are expected to be valid for long times (*t* → ∞), large sizes (*n* → ∞), and slow fragmentation (*β/r* < 1).

| | $\nu_F = 1$ | $\nu_F = 2/3$ | $\nu_F = 0$ |
| --- | --- | --- | --- |
| distribution exponent, $\mu$ | $1 + \frac{1}{1-\beta/r}$ | $\frac{5}{3} + \frac{\beta}{r}$ | $1 + \frac{\beta/r}{1+\beta/r}$ |

### Model summary

Clusters grow according to

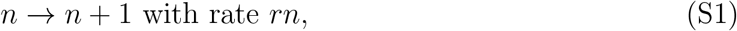

and they fragment according to

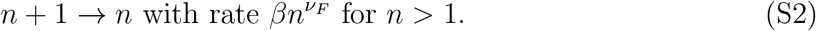

The cell lost during fragmentation becomes its own cluster of size one.

We now consider the deterministic dynamics of a large system. Putting these reactions together, we can write the master equation for the density of cells of size *n, c_n_*:

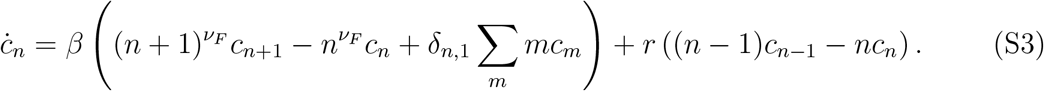

In what follows we will use the terms “density” and “total number” interchangeably, measuring volume in units of our system size (i.e., number of cells per gut). The first moment of this equation gives the total number of cells,

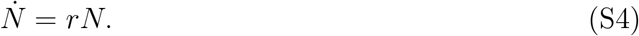

The zeroth moment gives the total number of clusters,

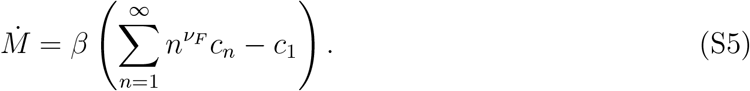

Here, the *c*_1_ term reflects the fact that in this model, cells must have size 2 or greater to fragment.

Finally, in a continuum picture, the size of a particular cluster is described by

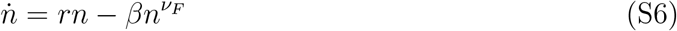

A well-known heuristic derivation of the stationary distribution of this type of process is based on the relationship between the number of clusters, *M* = ∑*_n_ c_n_*, and the total number of cells, *N* = ∑*_n_ nc_n_*. The key to this derivation is to recognize that *M*(*t*) acts as a proxy for the rank of the cluster that arises at time *t*: for the *j^th^* cluster to arise, there are *j* – 1 clusters that have a larger size, since the relative ordering of cluster sizes is preserved during exponential growth. For large sizes, when cluster rank is expressed as a function of cluster size it becomes proportional to the reverse cumulative distribution function, from which we obtain the density.

It turns out that the differences in behaviors of exponents measured in simulations for different values of *ν_F_* can be understood by considering the importance of two terms in particular: the *c*_1_ term in the equation for *M*, and the *βn^ν_F_^* term in the equation for *n*.

#### Case 1: *ν_F_* = 1

The total number of cells follows simple exponential growth, *N*(*t*) ~ exp(*rt*). For *ν_F_* = 1, the total number of clusters is governed by the equation

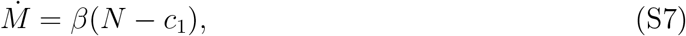

where the *c*_1_ term arises in our model because clusters can only fragment if they have size ≥ 2. At long times, however, we expect *c*_1_ ≪ *N* and we therefore ignore this term, leading to *M*(*t*) ~ (*β/r*)exp(*rt*) ~ *N*(*t*). A cluster that arises at time *t*’ will at a later time *t* have a size

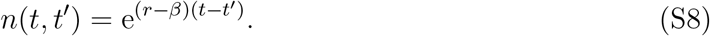

Ignoring overall *t* dependence, we can express this size as a function of the rank of this cluster,

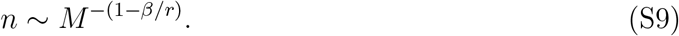

Inverting this relationship, and invoking the proportionality between *M* and the reverse cumulative distribution, *P*(size > *n*), results in

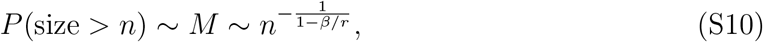

and differentiating produces the expected result

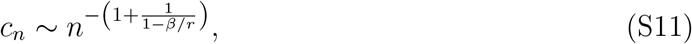

where *c_n_* is normalized by ∑ *c_n_* = *M*. This result matches the traditional Yule-Simons process, where each organism divides at rate *r* and then mutates with probability *ϵ*, with *ϵ* = *β/r*.

#### Case 2: 0 < *ν_F_* < 1

In this case, when considering the equation for the size of a particular cluster,

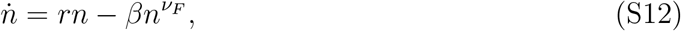

for *ν_F_* < 1 we ignore the second term on the right hand side. This term represents loss due to fragmentation, can be ignored for large sizes. Specifically, we consider sizes greater than a critical size, *n_c_* = (*β/r*)^1/(1−*ν_F_*)^ below which clusters will shrink. Ignoring this term, the size of a particular clusters that arose at time *t*’ is given by

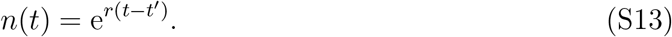

The total number of clusters follows

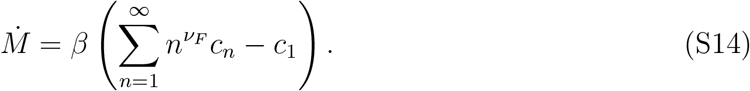

Like above with *ν_F_* = 1, we ignore the *c*_1_ term. Unlike for *ν_F_* = 1, we don’t have a closed equation for the fractional moment 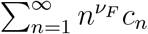. Therefore, we take the approach of making a power law ansatz

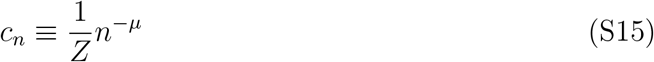

with normalization

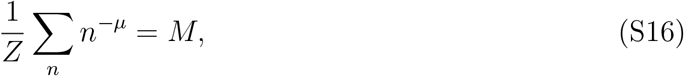

and then solve for the exponent *μ* self-consitently First, we approximate the sums by integrals and arrive at an equation for *M*

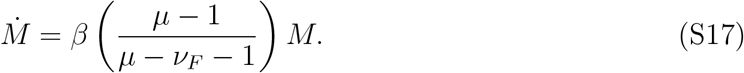

Then we follow the same logic as for the *ν_F_* = 1 case. Solving for *M*(*t*), we get

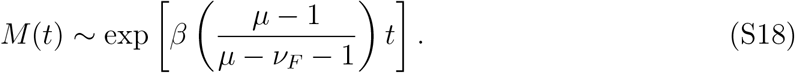

Combining terms into

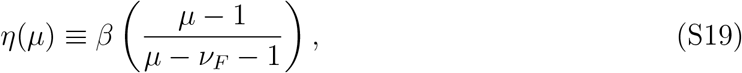

we then relate the size of a cluster that arose at time *t*’ to the rank of that cluster, *M*(*t*’),

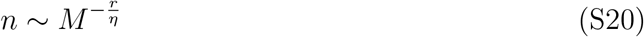

from which we compute the scaling behavior of *c_n_*,

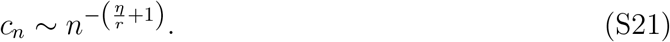

Equating this expression with the original ansatz, we arrive at a self-consistency equation for *μ*

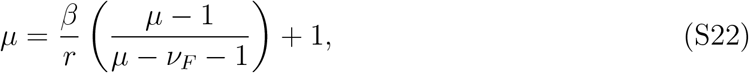

which we solve to obtain an exponent linear in the rates,

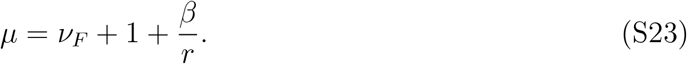

This result is plotted in Fig. 3D of the main text with *ν_F_* = 2/3 and agrees reasonably well with simulations.

#### Case 3: *ν_F_* = 0

For *ν_F_* = 0, the equation for *M* simplifies to

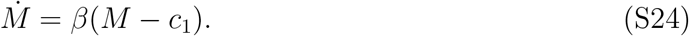

The simulation results in Fig. 3D indicate that the relationship between *μ* and *β/r* is no longer linear, which we expect to be due to the *c*_1_ term reducing the propensity for fragmentation. That this single-cell effect is relevant for *ν_F_* = 0 makes sense because we expect most clusters to be of order 1, which would make *c*_1_ of order *M*. To account for this term explicitly, we make the same power law ansatz as before,

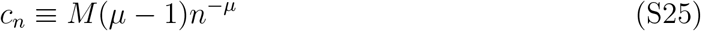

and then extrapolate down to *n* = 1 to estimate *c*_1_,

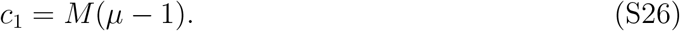

This extrapolation is purely a convenient approximation, as the distribution is likely not a true power law down to sizes of 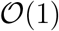. With this ansatz, the equation for *M* reads

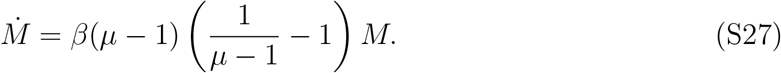

Combining terms we can define

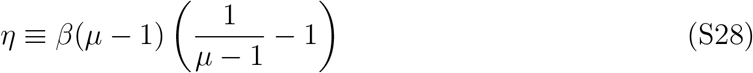

and write

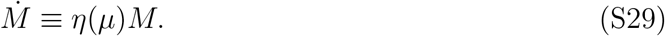

Then, following the same protocol as above, we can relate the frequency of a cluster to its rank,

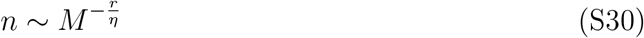

from which we compute the scaling behavior of *c_n_*,

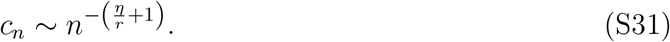

Equating this result with the original ansatz, we get the self-consistency equation

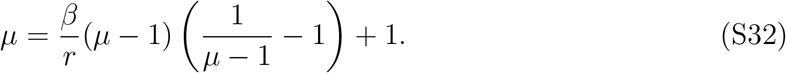

Solving this equation we get

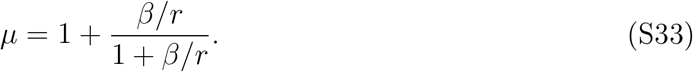

This result is plotted in Fig. 3D of the main text and agrees reasonably well with simulations, with notable deviations occurring once *β/r* ≈ 1.

### Discussion

In this model growth and fragmentation are treated as separate processes. This choice is convenient in the context of the full model including aggregation because classic reversible aggregation models [32] are contained within this general framework when growth and expulsion rates are set to zero, and also because fragmentation conserves total cell number. As a consequence of this choice, single cells are forbidden from fragmenting and don’t contribute to the total rate of fragmentation events. This feature differs from common evolutionary variants of this model for asexual populations, where growth and mutation are linked. In those models, all organisms divide and in each division have a probability of mutating (analogous to fragmenting in our model), so the rate of mutant production scales with the total population, including singletons. This is mostly a minor difference, but, as we showed, it does lead to different behaviors of the resulting distribution exponents in the limit of fast fragmentation.

We showed that we can account for this effect when it is important (in the case *ν_F_* = 0), but we cannot say *when* it will be important. That is because this effect depends on the number of single cells, which lies outside the regime of our large-size asymptotics that underly the continuum approximation. Ultimately the effect depends on how large the the number of single cells, *c*_1_, is compared to the fractional moment ∑*_n_ n^*ν_F_*^c_n_*. If the distribution has a significant shoulder, than c1 may be smaller than the extrapolation of the power-law form down to *n* = 1. In that case, the single-cell effect may be less important than this extrapolation would predict it to be.

## Supplementary Figures

**Figure S1:**
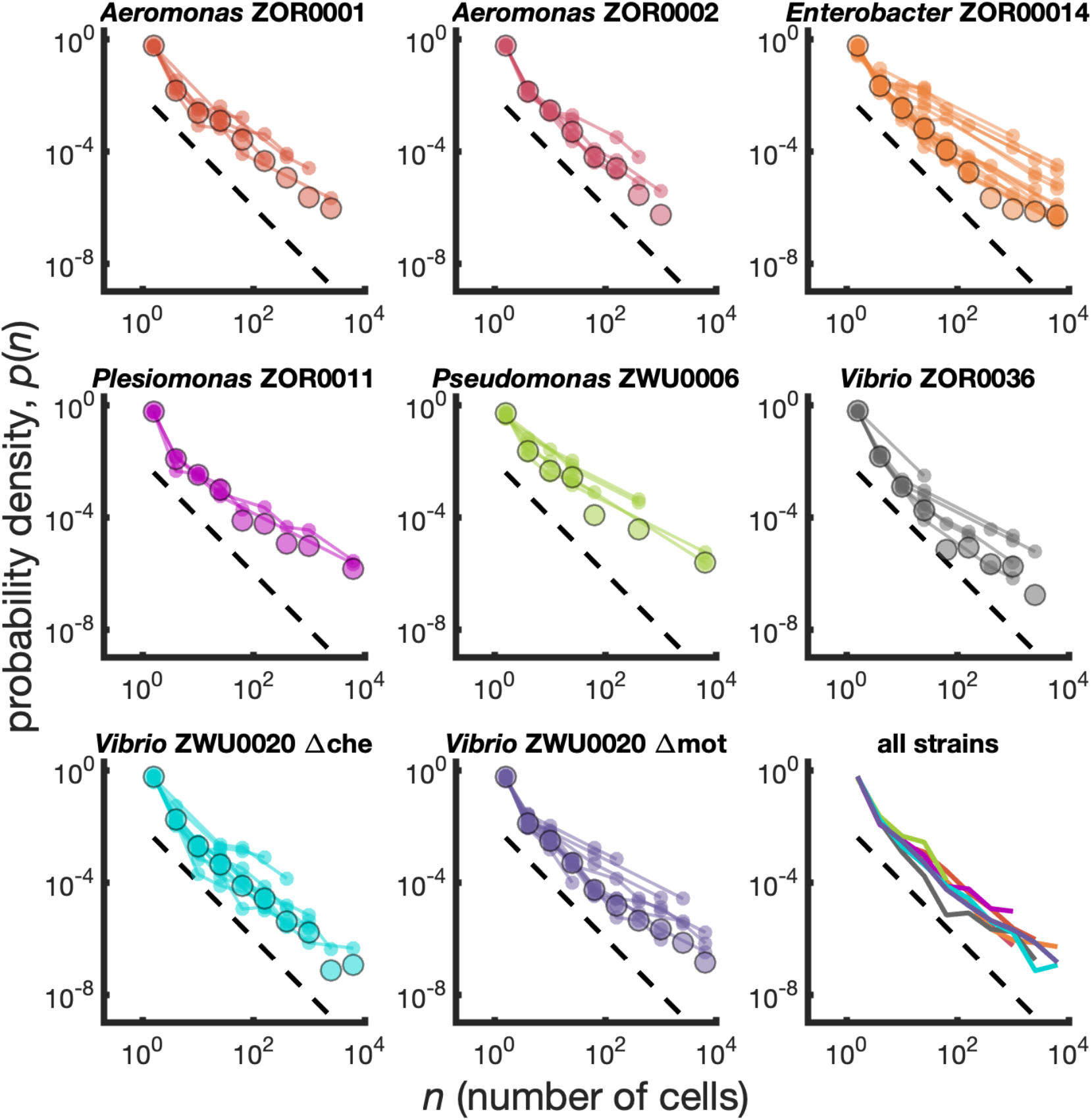
Different bacterial species exhibit similar cluster size distributions. Probability densities for eight bacterial strains monoassociated in larval zebrafish intestines. Small circles connected by lines represent the distributions constructed for each fish. Large circles are the result of pooling together sizes from all fish. Dashed line represents *p*(*n*) ~ *n*^−2^. Bottom right panel shows the pooled distributions for each strain as solid lines. Summary of data is given in Table 1.

**Figure S2:**
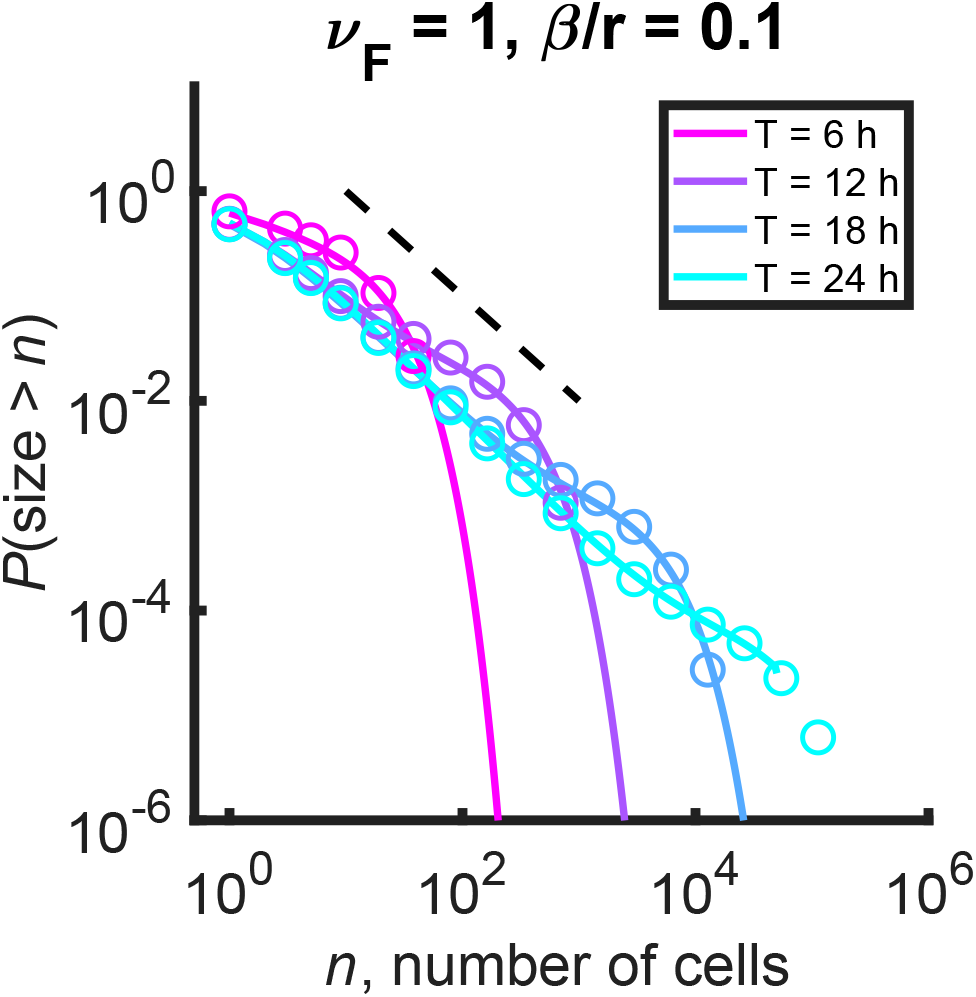
Mild curvature appears in the minimal growth/fragmentation process distribution at finite time, but the result is inconsistent with experimental data. Circles are the result of stochastic simulation, solid lines are the result of numerical integration of the master equation. Color denotes simulation time. Dashed black line: *P*(size > *n*) ~ *n*^−1^.

**Figure S3:**
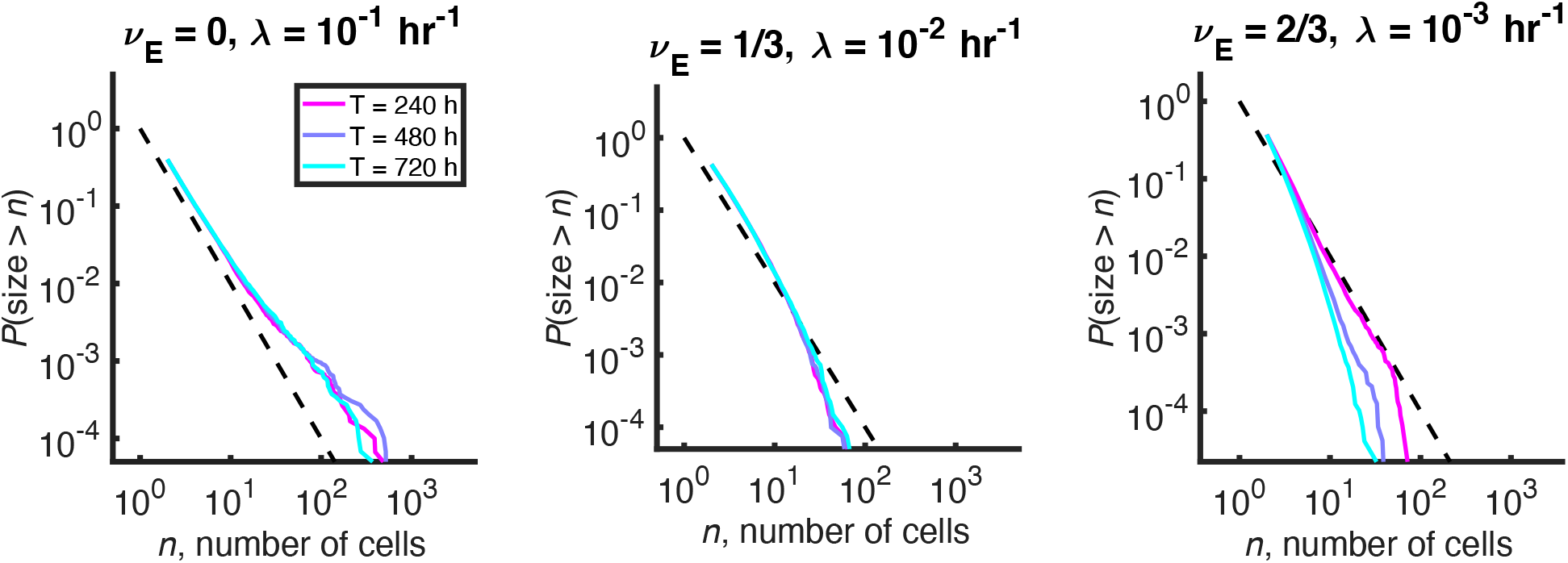
A modified process with carrying capacity and expulsion does not produce a plateau in the stationary size distribution. Reverse cumulative distributions computed from stochastic simulations with different values of the expulsion exponent, *ν_E_*. For each value of *ν_E_*, the expulsion rate, λ, was chosen such that for clusters of size *K* = 1000 cells, λ*Kν_E_*= 0.1 hr^−1^, consistent with experimental data. Three different simulation times are shown in each panel in differently colored solid lines. Long simulation times are required to approach steady state when λ becomes small, and the steady state is not quite reached in the right panel even after 720 hours. Dashed line indicates *P*(size > *n*) ~ *n*^−1^. Other parameters: *r* = 0.5 hr^−1^, *ν_F_* = 2/3, *β* = 0.5 hr^−1^, and the number of simulations = 100.

**Figure S4:**
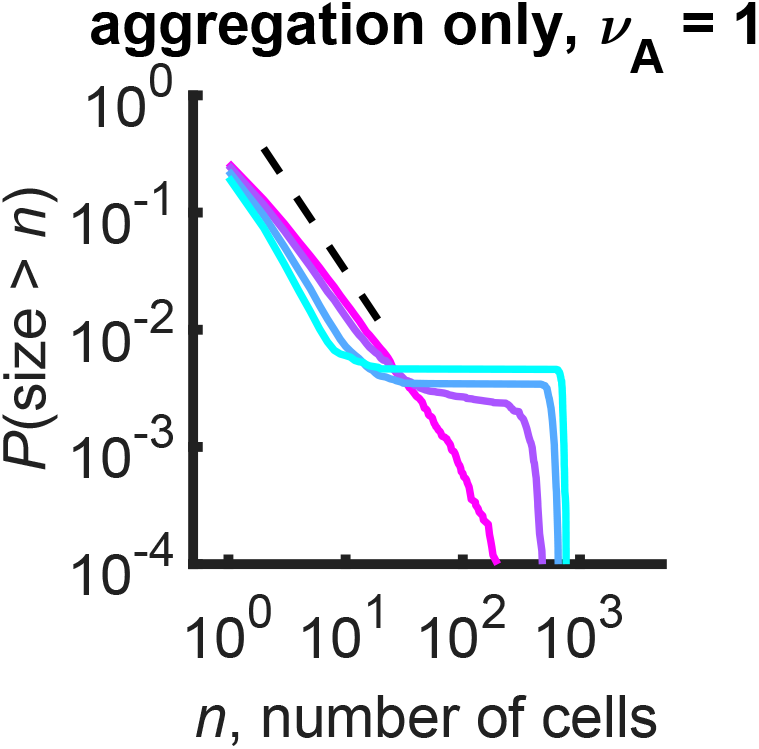
Plateaus arise at the gelation transition of purely aggregating systems. Reverse cumulative distributions computed from stochastic simulations are shown. Different curves represent different simulation times, ranging linearly from 0.1 to 0.175 hr (magenta to cyan). Dashed line represents an approximate analytic prediction of *P*(size > *n*) ~ *n*^−3/2^ in the sol phase at the transition point. Parameters: *α* = 0.01 hr^−1^, number of cells = 10^3^, number of simulations = 100.

